# Widespread Genotype-Phenotype Correlations in Intellectual Disability

**DOI:** 10.1101/220608

**Authors:** Emily L. Casanova, Zachary Gerstner, Julia L. Sharp, Manuel F. Casanova, F. Alex Feltus

## Abstract

**Background:** Linking genotype to phenotype is a major aim of genetics research, yet many complex conditions continue to hide their underlying biochemical mechanisms. Recent research provides evidence that relevant gene-phenotype associations are discoverable in the study of intellectual disability (ID). Here we expand on that work, identifying distinctive gene interaction modules with unique enrichment patterns reflective of associated clinical features in ID.

**Methods:** Two hundred twelve forms of monogenic ID were curated according to comorbidities with autism and epilepsy. These groups were further subdivided according to secondary clinical symptoms of complex versus simple facial dysmorphia and neurodegenerative-like features due to their clinical prominence, modest symptom overlap, and probable etiological divergence. An aggregate gene interaction ID network for these phenotype subgroups was discovered using via a public database of known gene interactions: protein-protein, genetic, and mRNA coexpression. Additional annotation resources (Gene Ontology, Human Phenotype Ontology, TRANSFAC/JASPAR, and KEGG/WikiPathways) were utilized to assess functional and phenotypic enrichment modules within the full ID network.

**Results:** Phenotypic analysis revealed high rates of complex facial dysmorphia in ID with comorbid autism. In contrast, neurodegenerative-like features were overrepresented in ID with epilepsy. Network analysis subsequently showed that gene groups divided according to clinical features of interest resulted in distinctive interaction clusters, with unique functional enrichments according to module.

**Conclusions:** These data suggest that specific comorbid and secondary clinical features in ID are predictive of underlying genotype. In summary, ID form unique clusters, which are comprised of individual conditions with remarkable genotypic and phenotypic overlap.

## BACKGROUND

Phenomics is a new and emerging area of study, underlying the development of genotype-phenotype mapping and the identification of different disease interaction networks [1]. Intellectual disability (ID) is a complex and highly heterogeneous group of disorders despite cognitive and behavioral overlap. Genotype-phenotype correlations have been reported within individual syndromes and across different mutations within a given gene, yet only recently have there been reports of more extensive genotype-phenotype clusters comprising subsets of the condition [2].

Previously, we reported associations between autism and epilepsy comorbidities in monogenic ID with trends in functional gene enrichment, suggesting these behavioral/neurological phenotypes represent etiological divergence at the molecular level [3]. Within these ID, additional secondary clinical features are also prominent, such as multiple congenital anomalies (MCA), neurodegeneration, brain atrophy, and motor disorders like upper motor neuron disease (UMND), all of which co-vary to greater or lesser degrees. Because of the prominence of these secondary clinical features, we have elected to extend similar work as Kochinke et al. [2] in order to perform in depth investigation into gene functional and modular enrichment in association with these features, in the hopes that in using a more general approach across an array of different disorders we may identify previously unseen genotype-phenotype associations.

In this study, we have found multiple unique gene clusters with specific functional enrichments that coincide with distinctive clinical phenotypes, indicating ID genes do indeed exhibit associations with phenotype. These broad associations may therefore help advance our knowledge of these disorders, as well as begin to delineate clusters of rare syndromes that were once thought unrelated.

## METHODS

### Gene-Phenotype Curation

Our gene-ID dataset was curated as described in [3]. To summarize the curation process, a comprehensive list of different forms of ID with known molecular origins was accessed from the Mendelian Inheritance in Man (MIM) database [4]. By selecting conditions with ID, we were able to estimate genetic penetrance for the autism and epilepsy phenotypes according to rates of comorbidity. Keywords for initial accession included: “intellectual disability,” “mental retardation,” “mentally retarded,” “global developmental delay,” “severe developmental delay,” and “profound developmental delay.” Any rare conditions not accessed by these call words were not included in the study for the sake of consistency. In addition, conditions were removed if they fulfilled any of the following criteria: 1) the ID was variably expressed and not considered a primary feature; 2) onset of ID was later than three years of age; 3) the condition was often lethal in infancy or early childhood; 4) the condition was considered genetically complex (e.g., deletion/duplication syndromes), with the exception of chromosome 2p16.3 deletion syndrome, which contains only the *NRXN1* gene; 5) autism was a defining symptom for diagnosis, as in the case of certain “susceptibility” genes; 6) the condition had < 2 reported cases; 7) the condition was a chromosomal instability syndrome, leading to an accumulation of different mutations; and 8) the condition was demarcated by a “?” indicating an unconfirmed or potentially spurious mapping.

The larger group was then subdivided according to comorbidities with autism and epilepsy and their frequencies, which were verified both through MIM and the larger literature (email ELC for dataset). For this study, only conditions with high autism and/or epilepsy rates, or without either comorbidity, were retained.

In addition, all conditions that were not contained within MIM’s Clinical Synopses were removed, resulting in a final dataset of 212 different conditions. The autism group with/without epilepsy (referred to here as the “autism group”) contained 59 unique conditions; ID with epilepsy but without autism (referred to as the “epilepsy group”) was composed of 83 unique conditions; and ID without autism or epilepsy (ID group) was composed of 70 unique conditions. (Email ELC for dataset.)

Comorbidity frequencies between ID and autism/epilepsy were obtained from the literature and described in detail in [3]. (Email ELC for additional data.) A high cut-off for inclusion within both the autism and epilepsy groups was ≥ 20% for all conditions for the sake of relative homogeneity. Only conditions without any indications of autism or epilepsy comorbidities, including the exclusion of single case examples, were placed within the ID group. This curation process culminated in three groups of conditions with very distinctive clinical and genetic profiles, as will be discussed in the Results section.

All conditions were annotated using the MIM’s Clinical Synopses (12/15/2016), which represent common clinical features of a disorder and are organized anatomically. According to [5], features included within Clinical Synopses:

> *… are taken from the literature and incorporated into the synopsis using a semi-controlled vocabulary. Many features include modifiers and additional terminology specific to medical subspecialties that are helpful for delineating overlapping disorders and distinguishing characteristic features. **Among genetically heterogeneous disorders, care is taken to include only those features that are present in patients with mutations in the same causative gene** [our emphasis].*

Conditions were annotated according to the presence of congenital anomalies in the following organs/tissues: the facial suite (face, eyes, ears, nose, mouth, dentition, neck); the cranial suite (cranial volume, synostoses, other cranial malformations, e.g., bitemporal narrowing); hands and feet; the limbs; the viscera and genitals (changes to the latter not otherwise due to peripubertal hypogonadism, etc.); hair and skin; and the brain (partial/complete agenesis of the corpus callosum and malformations of cortical development (MCD), the limbic system, the midbrain, and the brainstem, all visible via MRI). Complex (CFD) and simple facial dysmorphia (SFD) were annotated according to the number of facial regions affected, rather than according to the number of specific dysmorphisms associated with a given condition. Tissue regions include overall facial shape; the nose; the exterior of the mouth; the interior mouth such as tongue, dentition, and jaw shape; the form of the eyes; the midface (cheeks); and the ears. CFD was defined according to three or more malformations in distinct tissue regions, while SFD was defined as 1-2.

Phenotype interactions were analyzed across all congenital anomalies. Following analysis (see Results), CFD was selected as a defining secondary clinical feature for further genetic study, due both to clinical prominence and predictive ability in the presence of MCA syndromes. SFD were also selected as a secondary feature of interest for the sake of contrast, although were generally not predictive of MCA syndromes.

Conditions were also annotated for the presence of: neurodegeneration (confirmed according to literature search); brain atrophy; symptoms indicative of UMND, such as spasticity and hyperreflexia; and the presence of symptoms indicating the co-occurrence of 2 or more distinct movement disorders (UMND, lower motor neuron disease [LMND], disorders of the cerebellum, and disorders of the basal ganglia). Because brain atrophy and motor disorders were positively associated with neurodegeneration (see Results), all of the above clinical features were collapsed into a single category, “neurodegenerative-like features (NLF),” for the purposes of further genetic study. CFD and NLF phenotypes were further substantiated using the Human Phenotype Ontology (HPO) database, and, when that was insufficient, the general literature in order to ensure reliability of MIM’s Clinical Synopsis results for each of the conditions studied [6].

In order to study the association of the above clinical phenotypes with autism, epilepsy, and ID groups, conditions were subdivided according to the overlapping clinical phenotypes presented in Figure 2. This resulted in 18 unique gene sets, composed of 216 genes representing 212 different forms of monogenic ID (email ELC for dataset).

### Extended Gene Interaction Network

The GeneMANIA gene interaction database (genemania.org; [7]) was queried to discover additional known interactions for all 216 curated seed genes (email ELC for dataset). The database provides a report containing several different interaction types including physical, genetic, pathway, predicted, co-localization, co-expression, and shared protein domains. All interactions were recorded, but for the purpose of this study only genes with physical, genetic, or co-expression interactions were included in the finalized network (email ELC for dataset).

Visualization and analysis of the network was conducted via Cytoscape [8]. The Network Analysis function for undirected graphs was used to describe attributes including centrality, average connectivity, and clustering co-efficient using default parameters. The clusterMaker MCL algorithm was used to discover 20 gene clusters in the extended network (http://www.cgl.ucsf.edu/cytoscape/cluster/clusterMaker.shtml). Kochinke et al. [2] reported that nearly half of all ID genes physically interact with one another, with more than a third forming a single large interactive network. Therefore, we tested if phenotype labels, assigned at the gene curation stage, and their extended interactions were non-randomly assigned to MCL clusters.

In addition, because there is a portion of genes within the autism gene group that are not currently contained within the syndromic category of the SFARI gene database and may therefore be suspect, we have also assessed nonrandom clustering of syndromic SFARI seed genes to illustrate that similar clustering still occurs with more stringent exclusion criteria. Our approach was identical as in the main network analysis, with the exception that only seed genes contained within the syndromic SFARI category were used [9].

Enrichr was used for functional enrichment analysis on the main gene network, a platform that provides the following libraries used in this study: Gene Ontology (GO), KEGG/WikiPathways, TRANSFAC/JASPAR Position Weight Matrix (PWM), MGI Mammalian Phenotype (MP), and Human Phenotype Ontology (HPO) [10]. The Enrichr platform provides adjusted p-values using the Enricher list randomization method [11], Z-scores, and combined scores for each item. We used an adjusted significance threshold of *p* < 0.05.

### Statistical Analyses

For phenotype analyses, between- and within-group comparisons were performed using two-sample proportion Chi-square tests with a false discovery rate *p*-value adjustment (R pairwise.prop.test). Odds ratios with sample size adjustments [12] were computed to examine associations amongst different congenital anomalies, as well as associations within NLF, the latter without sample size adjustment.

For the purposes of assessing robustness of non-random assignment of seed and interactive genes to their respective MCL clusters, a power law distribution was calculated.

## RESULTS

Although all of the conditions reported here are rare, the average number of available cases ranged from 11-23 cases (*SD* = 13-25) across each of our three primary groups: autism, epilepsy, and ID only. There was a modest group difference (*χ ^2^* = 8.948, *p* = 0.0114), however this result was largely driven by comparison of the autism vs. epilepsy groups (*p* = 0.009), the former which presented with case numbers as high as 130. These results suggest that the number of available cases was likely an adequate minimum for our intended purposes.

We also studied the severity of ID across our main groups, in order to address concerns that autism and epilepsy may be related to the level of cognitive impairment rather than molecular etiology. Referencing the literature, we annotated a range of severity (1 = mild, 2 = moderate, 3 = severe, 4 = profound) for each condition and selected the median as a representative value. We find that there are indeed between-group differences (*p* = 0.001) but this result is primarily driven by a comparison of the epilepsy (*Med* = 2.54, moderate-to-severe) and ID groups (*Med* = 2.06, moderate) (*p* = 0.001). Meanwhile, the autism group (*Med* = 2.31, moderate-to-severe) does not significantly differ from epilepsy or ID (*p* = 0.0742). This suggests that ID severity does not differ dramatically between our three groups, although the ID only group exhibits a broader range than its autism and epilepsy counterparts (Fig. 1A).

**Figure 1.**
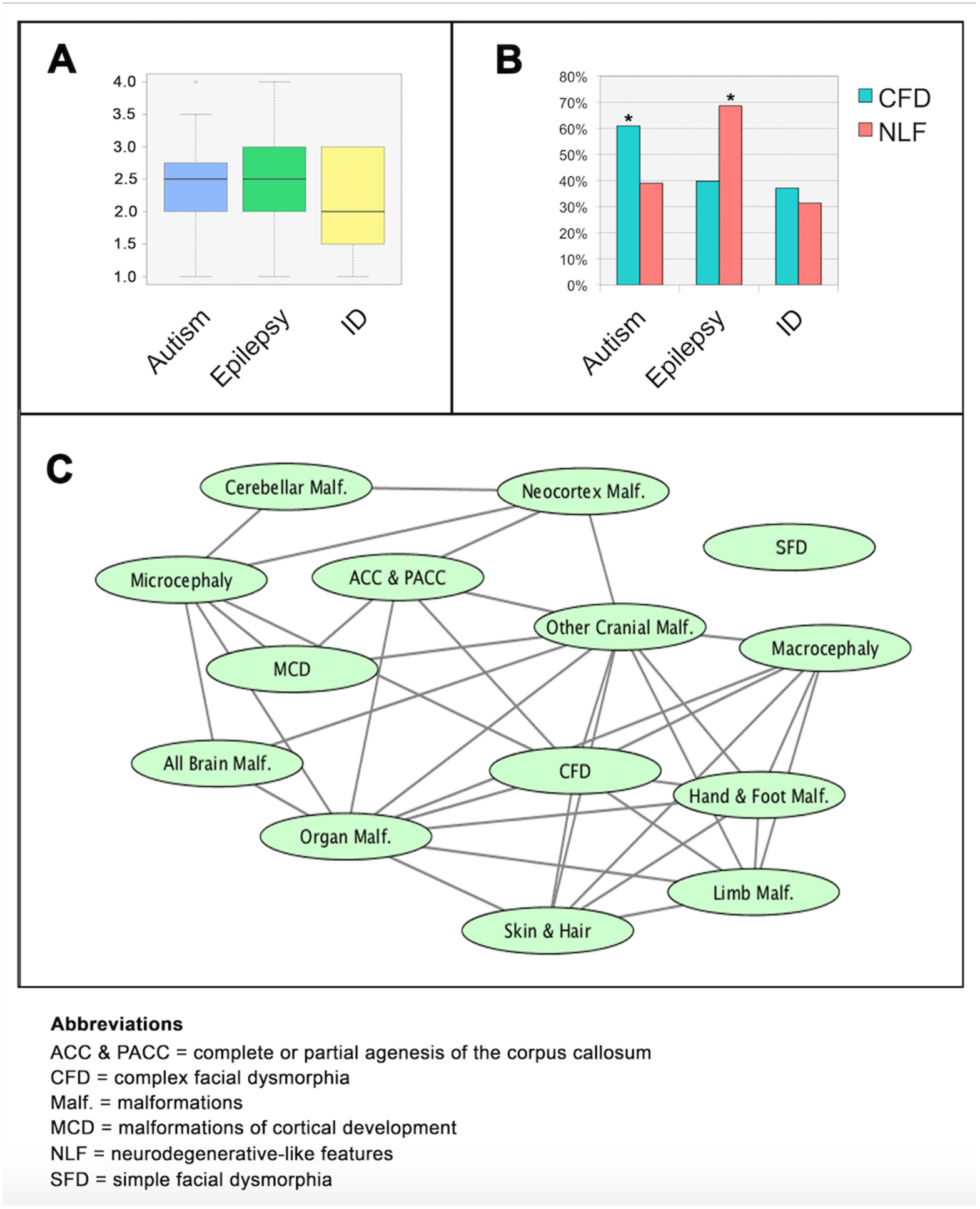
***Phenotype Data.**(A) Range of severity of intellectual disability (ID) across the three main groups. Y column values: 1 = mild, 2 = moderate, 3 = severe, 4 = profound. Autism and epilepsy share the greatest similarity, while ID only differs significantly from the epilepsy group presenting with broader range of cognitive phenotypes. (B) Occurrence of complex craniofacial dysmorphisms (CFD) and neurodegenerative-like features (NLF) across the three main groups. (See also Fig. 2.) (C) Association network between the major physical dysmorphia. Edges indicate a significant association between two given features.*

**Figure 2.**
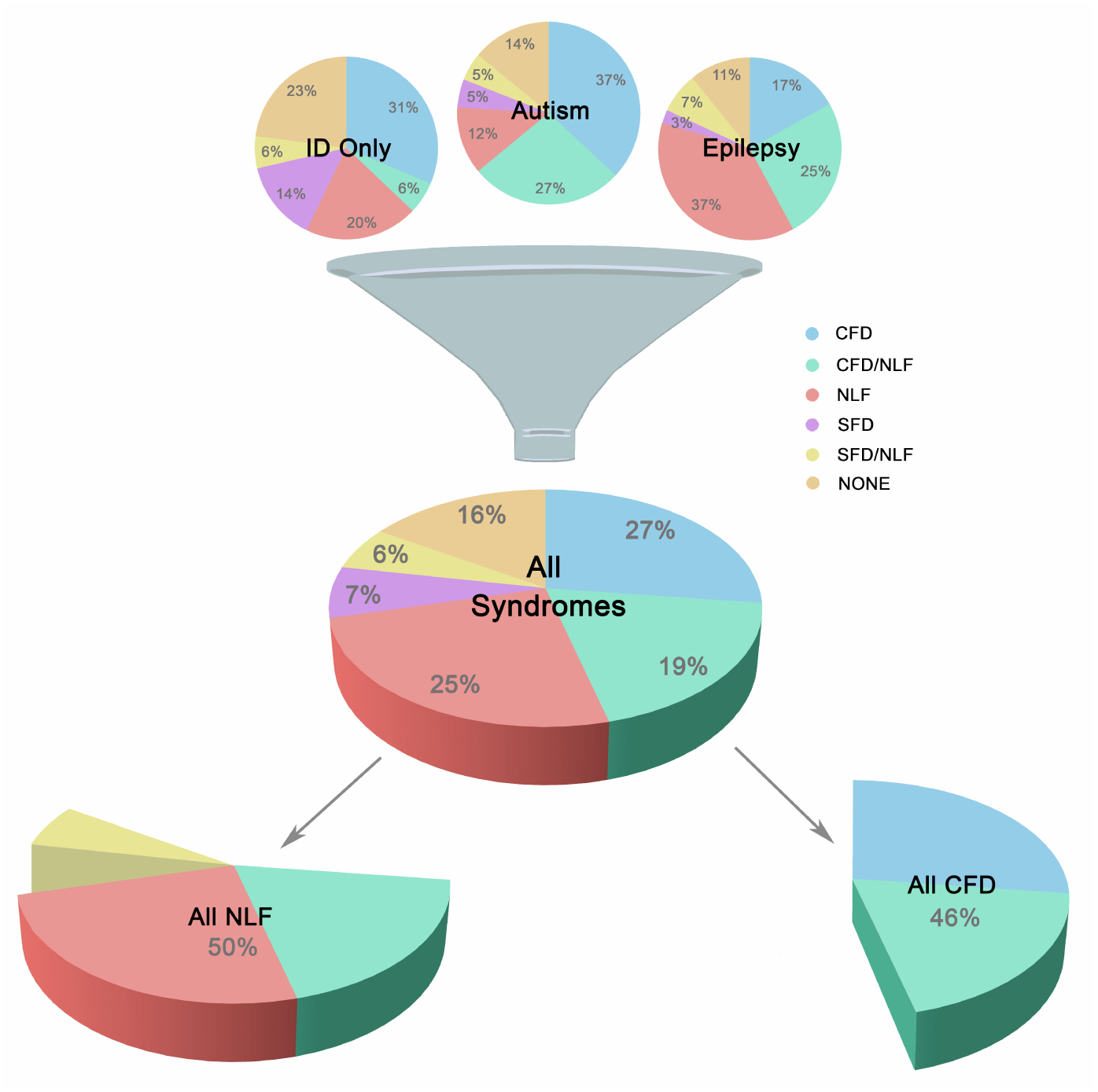
***Major Subgroups.** Illustration showing breakdown of the three main groups (autism, epilepsy, and ID only) into smaller clinically-related subgroups. Subgroups were defined according to the presence of complex facial dysmorphia (CFD), neurodegenerative-like features (NLF), simple facial dysmorphia (SFD), or a lack of these same features (i.e., “none”). As shown in the lower portion of the image, CFD and NLF overlapped approximately 20% of the time, although analyses indicate no clear relationship between the two sets of features, suggesting possible genetic pleiotropy when comorbid.*

### Clinical Features Common in Monogenic Intellectual Disability

Congenital anomalies are prominent features within monogenic forms of ID. In this study, the most common congenital anomaly reported was CFD, occurring in almost half of the conditions studied. Less frequent though still prominent congenital anomalies included (in order of frequency from most to least): microcephaly; organ malformations; brain malformations (visible via MRI); craniosynostoses and other cranial malformations; hand and foot malformations; skin and hair disturbances; SFD; macrocephaly; and limb malformations.

Not only was CFD the most common dysmorphism, it is also strongly associated with other types of dysmorphia (*z* = 0.7813-7.1947, *p* < 0.001-0.014; *OR* = 2.19-13.461; *OR CI* = 1.261-6.266, 3.805-28.717), with the exception of specific brain malformations (*z* = 0.328-2.230, *p* = 0.055-0.814; *OR* = 1.159-1.744; *95% CI* = 0.479-1.108, 2.654-4.892) (Fig. 1A). (Email ELC for dataset.) One primary exception was the strong relationship between complete/partial agenesis of the corpus callosum (ACC) and CFD, suggesting significant etiological links (*z* = 2.993, *p* = 0.009; *OR* = 4.465; *95% CI* = *95% CI* = 1.762, 15.117). Microcephaly was also only very weakly predictive of MCA (aside from brain and cranium) (*z* = 1.113-2.781, *p* = 0.014-0.369; *OR* = 1.393-2.19; *95% CI* = 0.734-1.261, 2.498-3.948), and therefore facial dysmorphia were annotated separately from deviations in cranial volume in this study, despite the clinical tradition of grouping all craniofacial malformations together.

Neurodegeneration was also common occurring in approximately 20% of ID and was an extremely strong predictive factor for the presence of brain atrophy and various movement disorders, especially UMND (*z* = 5.110, *p* < 0.001; *OR* = 8.61; *95% CI* = 3.77, 19.68) (email ELC for dataset). Another ∼30% of conditions displayed either brain atrophy, UMND, or multiple movement disorders (MMD) (or some combination thereof) but are not currently recognized as classical neurodegenerative disorders. However, because of their strong association suggesting linked etiologies, neurodegeneration, brain atrophy, UMND, and MMD were combined under a single heading, “neurodegenerative-like features” or “NLF,” for the purposes of this study (*z* = 4.69-8.73, *p* < 0.001; *OR* = 4.64-56.53; *95% CI* = 2.44-22.86, 8.82-139.82). NLF occurred in 50% of the conditions studied, overlapping CFD approximately 19% of the time. Despite this large overlap, in the majority of cases these features did not co-occur and, overall, exhibited no statistically significant relationship with one another (*p* = 0.515; *OR* = 0.834; *95% CI* = 0.483, 1.440). This suggests that while these phenotypes may co-occur in a large minority of these conditions, they are nevertheless unique symptom clusters and may instead reflect genetic pleiotropy when comorbid (see Fig. 2).

Previous results by Casanova et al. [3], utilizing a near-identical dataset, indicate a divergence in functional gene enrichment in ID according to autism and epilepsy comorbidities. Here we report additional clinical phenotype enrichment that varies according to these behavioral/neurological comorbidities. Namely, the autism group was significantly enriched for the presence of CFD (61% vs. 37-40%) (Fig. 1A), suggesting many rare autism syndromes may be dysplastic in nature (*χ^2^* = 5.42-6.38, *p* = 0.03) [13-15]. Meanwhile, the epilepsy group was similarly enriched for NLF (68% vs. 31-39%), indicating some form of cell stress involvement in these IDs (*χ^2^* = 11.18-19.63, *p* < 0.001) [16, 17]. There are additional clinical phenotypes that vary according to group, such as enrichment of neocortical malformations (identified by MRI) (*z* = 4.4566, *p* < 0.001, *OR* = 6.4289; *95% CI* = 2.836, 14.573) and microcephaly (*z* = 2.8656, *p* = 0.011, *OR* = 2.2778; *95% CI* = 1.297-4.000) in the epilepsy group. (Email ELC for dataset.)

### ID Genes Cluster According to Phenotype

Using a list of 216 seed genes divided according to our phenotypes of interest, we have identified an additional 354 interacting genes using the GeneMANIA gene interaction database (genemania.org). This resulted in the formation of 17 unique gene sets composed of a total of 1,195 genes upon which to perform cluster analyses according to protein-protein interaction (PPI), gene interaction, and mRNA co-expression. (One of the autism subgroups failed to show any significant intracluster interactions and therefore was not included in the cluster and functional enrichment analyses.)

As can be seen in Figure 3A, the seed genes plus PPI, gene-interacting, and co-expression loci form 17 sets of relatively non-overlapping clusters, constituting tight interaction/coexpression networks. Thirteen of the 17 gene sets form particularly tight clusters and are interconnected via specific hub nodes (Fig. 3B-E). (Email ELC for dull dataset.) Overall network degree distribution modestly fits the power law distribution (*r* = 0.776), indicating these networks trend towards scale free behavior. Other topological parameters of interest include: clustering coefficient = 0.342; centralization = 0.034; and average connectivity = 5.287. SFARI-only syndromic genes likewise formed similar non-random clusters (*r* = 0.687), indicating the robustness of the autism results overall (Fig. 3F).

**Figure 3.**
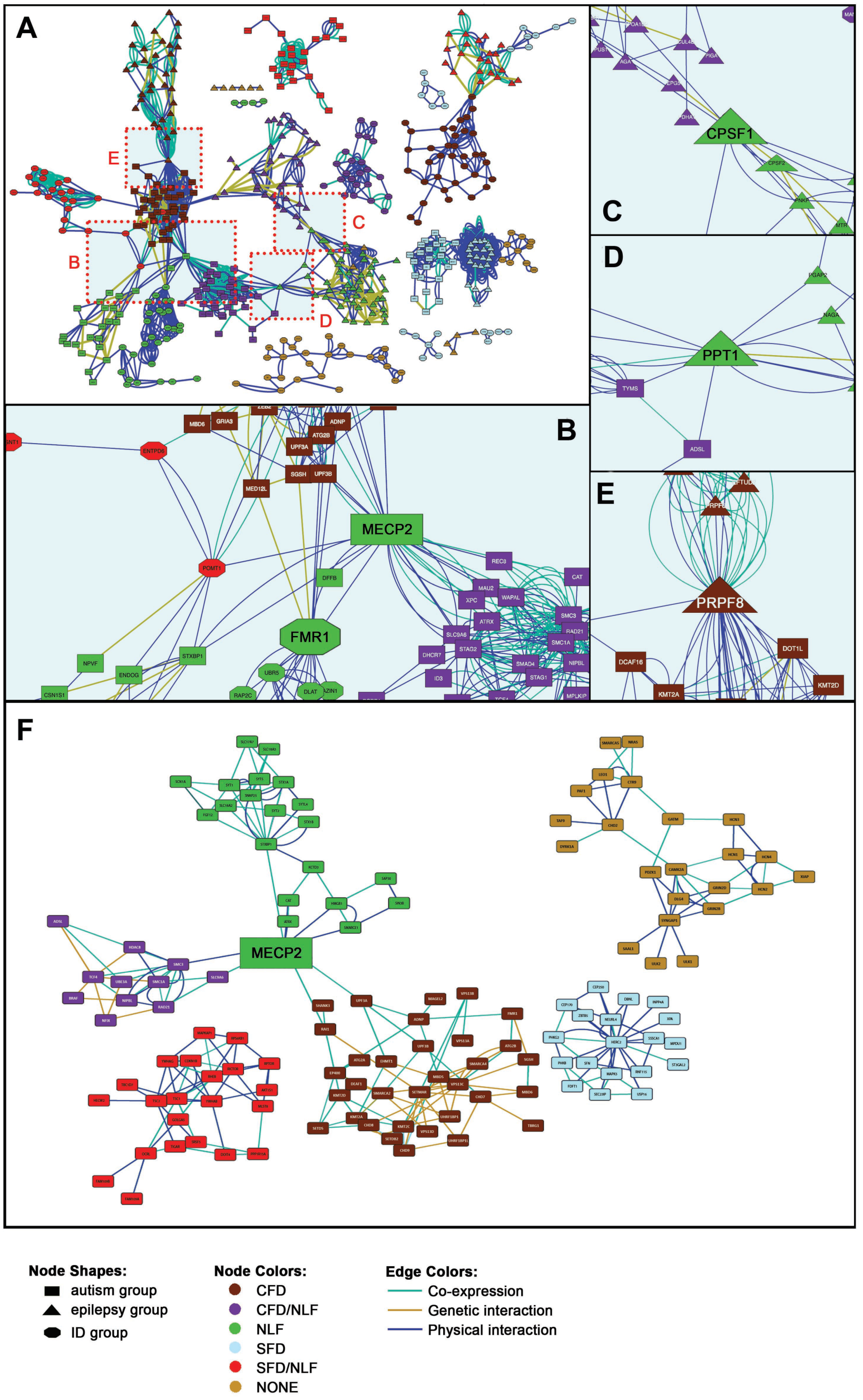
***Gene Interaction Network.** (A) Full gene interaction network. (Email ELC for detailed network image.) (B) Autism-linked MECP2 and FMR1 hubs. (C) Epilepsy-linked CPSF1 hub. (D) Epilepsy‐ and autism-linked PPT1 hub. (E) Epilepsy‐ and autism-linked PRPF8 hub. (F) Syndromic SFARI gene interaction network.*

Within the main network, more than half of the gene sets are interconnected via 10 hub nodes (Fig. 3B-E). The Rett Syndrome-associated gene, *MECP2,* for instance, forms a hub connecting half of the autism-related gene sets, particularly those with secondary clinical features of CFD, combined CFD/NLF, and pure NLF, as well as connecting one of the ID group clusters (Fig. 3B). In addition, *MECP2* remains an important hub node in the SFARI-only syndromic network, continuing to link CFD, CFD/NLF, and NLF autism subgroups. MECP2’s nature as a semi-ubiquitous repressor of long genes, which typifies many neural genes, places it in a key position to regulate development of the central nervous system and thus to potentially interact with many of the genes presented here [18].

Likewise, the Fragile X Syndrome-associated gene, *FMR1,* forms a major hub connecting the same clusters *as MECP2* within the main network, although this result is not maintained within the abbreviated SFARI network (Fig. 3B). Interestingly, like MECP2, there is some evidence to suggest that FMRP specifically targets gene products translated from long genes, suggesting MECP2 and FMRP may regulate different points along many of the same pathways [19, 20].

Two other major hubs in the main network are involved in mRNA processing: *CPSF1,* which is involved in 3’ processing of mRNA, and *PRPF8,* which acts as a scaffold for spliceosomal complexes and snRNA. As shown in Figure 2C, *CPSF1* connects two epilepsy modules with features of NLF; meanwhile, *PRPF8* (Fig. 2E) connects epilepsy/CFD (EPI/CFD) with autism/CFD (AUT/CFD). As we shall see in the following section, a number of the epilepsy clusters are enriched for mRNA processing. Interestingly, PRPF8 is also essential for sister chromatid cohesion, making it therefore surprising that it forms a hub with AUT/CFD rather than AUT/CFD/NLF, as we shall see in the following section [21].

Finally, the hub, *PPT1*, whose mutation is responsible for the neurodegenerative and lethal condition, Neuronal Ceroid Lipofuscinosis 1, links the EPI/NLF and AUT/CFD/NLF modules (Fig. 3D). As a glycoprotein involved in catabolism of lipid-modified proteins and a regulator of heat shock proteins, its loss results in excessive generation of reactive oxygen species (ROS) [22, 23]. *PPT1’s* role as a hub linking EPI/NLF and AUT/CFD/NLF can potentially be viewed in light of the roles chronic ROS play in the synaptic impairment that ultimately leads to a host of neurodegenerative disorders [24].

### Functional Enrichment Trends in Gene Modules

Some clusters showed little obvious trends in functional enrichment, such as EPI/SFD/NLF and ID/SFD. This may be a reflection of etiological diversity in these respective modules and/or the inadequacy of current platforms in estimating disparate functional relationships.

Other groups, however, appeared to show distinctive functional trends, particularly those associated with CFD. For instance, the AUT/CFD module is strongly enriched for processes relating to *chromatin modification (z =* -2.40, *p <* 0.001), *histone modification (z =* -2.39, *p <* 0.001), *methylation (z =* -2.45, *p =* 0.007), *transcription factor binding* (*z* = -2.16, p = 0.026), and is localized to the nucleus (*nucleolus*) (*z* = -2.21, *p* = 0.002) (Fig. 4A). More than a third of AUT/CFD genes are also transcriptional targets for Wilms tumor suppressor 1 (Wt1), a transcription factor that helps regulate cell development and survival (*z* = -1.62, *p* = 0.036). In addition, almost half of AUT/CFD genes are transcriptional targets of Lef1, a positive regulator of the canonical Wnt pathway, which is itself a foundational network involved in organ and tissue morphogenesis (*z* = -1.48, *p* = 0.036) [25].

**Figure 4.**
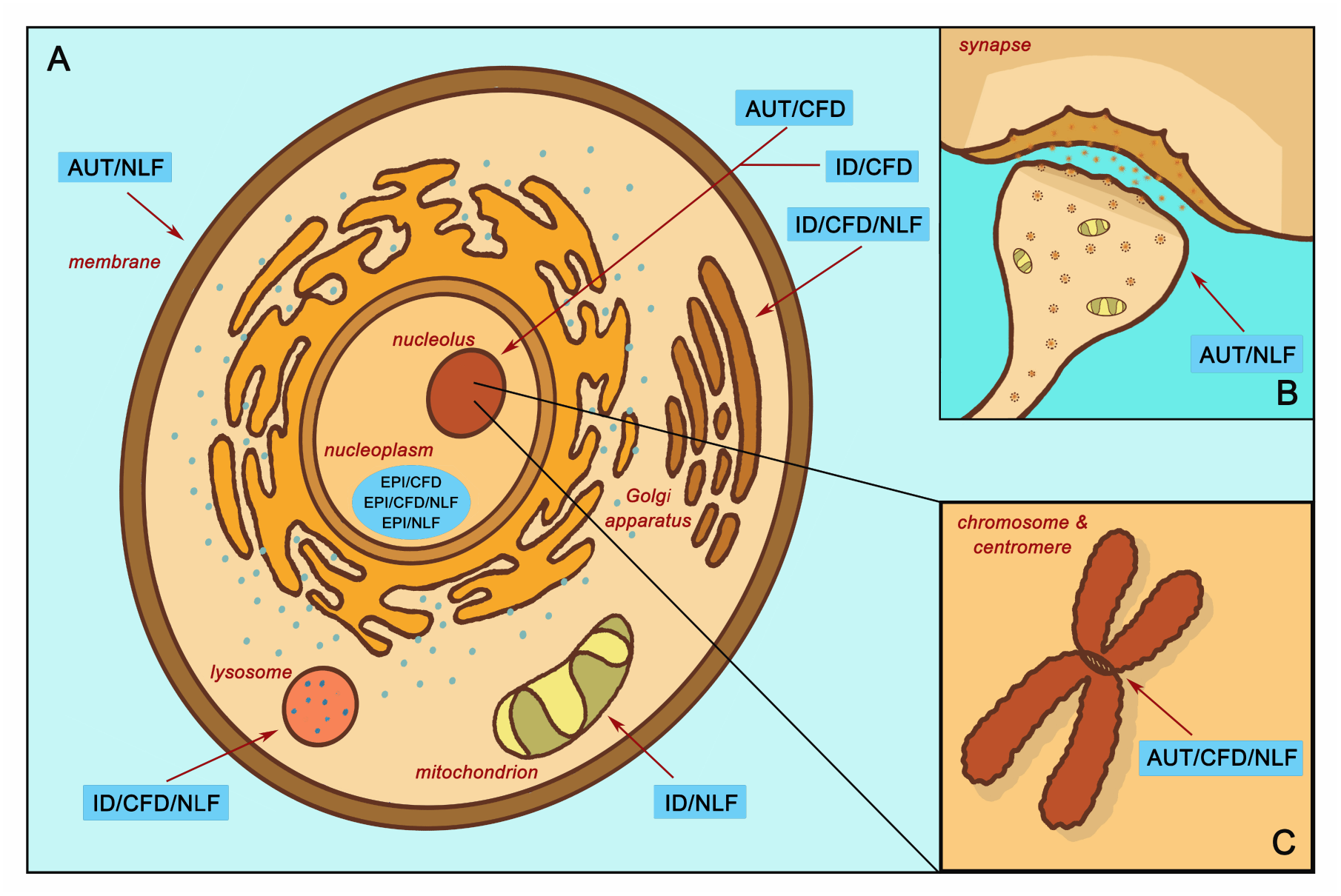
***Enrichment Localization According to Subgroup.**(A) Localized enrichment according to subgroups within the main body of the cell. (B) Subgroup enrichment within the synapse. (C) Subgroup enrichment at the chromosome and centromere. Abbreviations: intellectual disability with autism and with/without epilepsy (AUT); intellectual disability with epilepsy and without autism (EPI); intellectual disability without autism or epilepsy (ID); complex facial dysmorphia (CFD); neurodegenerative-like features (NLF).*

In contrast, the EPI/CFD gene cluster, though likewise relegated to the *nucleoplasm* (*z* = -2.16, *p* < 0.001) and involved in *histone modification* (*z* = -2.39, *p* < 0.001), is also enriched for processes involved in *mRNA processing* (*z* = -2.37, *p* = 0.003) and the *spliceosomal complex* (*z* = -2.15, *p* < 0.001). Similarly, EPI/CFD/NLF was enriched for *RNA polyadenylation* (*z* = -2.66, *p* <= 0.003). ID/CFD meanwhile is enriched in *kinase binding* (*z* = -2.55, *p* = 0.011) and *chromatin binding* (*z* = -2.45, *p* = 0.031), while ID/CFD/NLF is enriched for *protein glycosylation* (*z* = -2.34, *p* < 0.001) and is localized to the *Golgi membrane* (*z* = -2.29, *p* > 0.001) and the *lysosome* (*z* = -2.31, *p* > 0.001).

When comparing the two autism CFD modules to one another, we found that both AUT/CFD and AUT/CFD/NLF are involved in *chromatin binding* (*z* = -2.47, *p* < 0.001). However, AUT/CFD/NLF is also strongly enriched for processes involving the *mitotic cell cycle* (*z* = -2.30, *p* < 0.001) and *sister chromatid cohesion* (*z* = -2.67, *p* < 0.001), which is entirely missing from the AUT/CFD module.

In contrast to its CFD counterparts, ID/NLF was enriched in *hydrogen ion membrane transporter activity* (*z* = - 2.34, *p* = 0.003) and was involved in the *respiratory chain* (*z* = -2.59, *p* < 0.001) within mitochondria. In addition, it displayed pathway enrichment in relation to *Parkinson’s disease* (*z* = -1.77, *p* < 0.001), *Huntington’s disease* (*z* = -1.85, *p* = 0.002), and *Alzheimer’s disease* (*z* = -1.72, *p* = 0.015). The EPI/NLF module, in contrast, was enriched for a variety of terms, such as *myelin sheath* (*z* = -2.89, *p* < 0.001), *mRNA polyadenylation* (*p* = 0.007, *z* = -2.71), *carboxylic acid biosynthetic process* (*z* = -2.35, *p* = 0.007), and *protein folding* (*z* = -2.31, *p* = 0.007), suggesting that despite strong intracluster connectivity, the etiology of the EPI/NLF cluster is comparatively diverse. Meanwhile, AUT/NLF was modestly enriched for *membrane depolarization* (*z* = -2.26, *p* = 0.005), *regulation of postsynaptic membrane potential* (*z* = -2.09, *p* = 0.009), and *regulation of synaptic plasticity* (*z* = -2.15, *p* = 0.027). This indicates that disturbances to synaptic proteins in autism could be related to symptoms of NLF, an idea that may be worthy of further exploration in relation to autistic regression given the role of synaptic impairment in the etiologies of many neurodegenerative disorders [26]. Interestingly, recent research indicates that autistic individuals with gene disrupting mutations in postsynaptic density genes are more likely to experience autistic regression than individuals with mutations in genes of other functional classes [27].

## DISCUSSION

The present study provides evidence of genotype-phenotype correlations throughout multiple ID subsets. In particular, the presence of autism (with or without epilepsy), epilepsy (without autism), CFD, and NLF appear to be general predictors of associated gene function. The AUT/CFD module, for instance, is linked with genes localized to the nucleus. These genes are involved in chromatin modifications; histone modifications; methylation; transcription factor binding; and are key in regulating embryonic development. In contrast, gene products of the AUT/CFD/NLF cluster are likewise localized to the nucleus and involved in chromatin binding, but are typically involved in regulation of the cell cycle and sister chromatid segregation. Nuclear localization appears to be a strong risk factor in the developments of both autism and CFD in these clusters; however, cell cycle involvement may provide an additional risk for NLF as we see in many classical neurodegenerative diseases [28].

Nuclear localization, in general, seems to be a strong predictive factor for the presence of CFD and, more weakly, SFD, although specific functional enrichments vary with the presence of autism, epilepsy, and NLF accordingly. AUT/CFD, EPI/CFD, and ID/CFD all tend to be localized to the nucleus and are at least modestly enriched in processes relating to chromatin binding and modifications.

We have also identified a number of major hubs, linking otherwise non-overlapping gene modules. Although functional relevance of some of the hubs is currently uncertain, several of the autism hubs are already major foci within the current literature. Both *MECP2* and *FMR1*, for instance, have received considerable attention and *FMR1* in particular has been previously identified as a major pathway of interest in the pathophysiology of autism [29-31]. Our data therefore reinforce previous results and extend that data, indicating that *MECP2* and *FMR1* are foundational pathways in autism risk regardless of secondary clinical phenotype.

Finally, we have shown that specific secondary clinical phenotypes exhibit strong association with ID according to comorbidities with autism and epilepsy. For instance, the high rates of CFD and MCA in rare autism syndromes are strongly suggestive of a common biology despite genotypic variation. Despite the dearth of obvious brain malformations reported in our autism dataset, the high prevalence of microscopic dysplastic foci in idiopathic autism tends to validate this point [13, 15, 32, 33].

Our results have also shown that close to half of the conditions studied here exhibit features reminiscent of neurodegeneration, although only about a fifth are officially recognized as “neurodegenerative disorders.” The occurrence of NLF is particularly prominent in the epilepsy group, although functional enrichment of the ID/NLF subgroup is more aligned with processes of classic neurodegeneration. However, these data suggest that: 1) postmortem analysis of neurodegeneration may be understudied in some of these conditions, and/or 2) proteopathies with obvious inclusions may comprise only a subset of a broader range of neurodegenerative-like disorders, which have subtler, more complex etiologies with progressions that differ from the typical dementias that occur in later life. In support of this, Sarnat et al. [34] have recently addressed such concepts within the context of “infantile tauopathies,” such as tuberous sclerosis and focal cortical dysplasia 2. At present, recognized infantile proteopathies include solely those conditions resultant from MTOR overexpression, a known mechanism of neurodegeneration [35]. However, given the range of inclusion bodies associated with adult forms of neurodegeneration and senile dementias, the list of infantile proteopathies is likely to expand in future and may eventually be recognized as a major cause of some developmental and intellectual disabilities [34, 36].

### Current Limitations & Future Research

Given the nature of the MIM database, whose purpose is intended to summarize genetic and syndromic disease states, research procedures have varied across individual studies that compose the MIM. For these reasons, our results must be extrapolated cautiously, requiring further investigations at the clinical and molecular levels. However, in order to limit the extent of Type I errors, we have elected to study clinical phenotypes whose medical evaluations are standardized across health fields, ensuring that the clinical data reported here may be relatively reliable [37, 38].

One major exception to this is the field of autism diagnostics, which has changed significantly over the past 25 years. A majority (59%) of seed genes used in the cluster analysis is included within the syndromic category of the SFARI Gene Database, supporting their diagnostic reliability in this study. While we are unable to directly address diagnostic reliability of the remainder of autism genes, we instead assessed robustness of non-random clustering of this subset of syndromic SFARI genes, which like the larger autism gene group exhibited similar clustering. This supports our general findings as well as potential risk status of non-SFARI genes included in this study.

Another limitation of the study is the question of its applicability to a broader range of conditions. The study of severely affected individuals with rare genetic syndromes is a common approach to investigating human illness in order to better understand complex conditions. However, such assumptions are based on symptom similarity rather than biological evidence. As such, our results may not apply to forms of ID, autism, and epilepsy that lack strong genetic roots. However, recent work by Rossi et al. [39] suggest that even those patients with autism but without obvious syndromes often harbor potentially deleterious variants in many of the same genes studied here. Further lines of research will continue to address potential cross-applicability of the data presented here. In the meantime, we believe the clusters we’ve described can provide a platform for the further elucidation of common denominator pathways and the regulatory networks underlying these complex conditions, leading to the subtyping of disorders.

### Conclusions

The present study provides strong evidence that ID-associated phenotypes cluster according to related gene function. Specifically, gene modules form according to autism, epilepsy, CFD, and NLF comorbidities. Future research will help to delineate these clusters in greater detail, as well as determine whether additional genotype-phenotype correlations exist in these and related datasets.

## COMPETING INTERESTS

The authors declare that they have no competing interests.

## AUTHORS’ CONTRIBUTIONS

ELC and FAF conceived the study. ELC curated the phenotypic data and JLS performed statistical analyses on that data. ZG and FAF performed cluster and enrichment analyses and associated statistics. MFC provided expertise on autism, intellectual disability, and epilepsy and was integral in helping design the study as well as interpret results. All authors contributed substantially to the drafts and have read and approved the final manuscript.

## ACKNOWLEDGMENTS

This work was supported by the National Institutes of Health [grant number R01 HD-65279].

